# Multi-scale phenotyping of senescence-related changes in roots of *Rapeseed* in response to nitrogen deficiency

**DOI:** 10.1101/2024.04.03.587968

**Authors:** Maxence James, Céline Masclaux-Daubresse, Thierry Balliau, Anne Marmagne, Fabien Chardon, Jacques Trouverie, Philippe Etienne

## Abstract

Senescence related markers have been widely studied in leaves in many plant species. Root senescence is more difficult to characterize. The existence of two different root organs in *B. napus,* with a taproot that appear to be specifically dedicated to the storage and remobilization of nutrients, offered the possibility of analysing the temporality of the changes linked to aging, based on the degradation of the taproot reserves. Microscopic and biochemical analyses showed that taproot plays an important role in carbon and nitrogen storage as reflected by the large quantities of starch and proteins present at early development stages. The proteomic study associated to the description of biochemical, morphological and anatomic changes provides a comprehensive picture of the main events occurring in the taproot and in the lateral roots with aging. Master modifications as protein and cell wall degradation, amino acid catabolism versus synthesis, nucleic acid degradation are presented and senescence related markers specific or not of the root types were identified. Comparison with Arabidopsis public data facilitated the identification of markers common to root and leaf senescence. The analysis of protease changes provides a list of candidates that may play a role in nitrogen and carbohydrate remobilization from taproot to the shoot and flowering organs and that would deserve attention for further functional analyses.

## INTRODUCTION

Senescence corresponds to the final stage of the development of plant organs. It was mainly studied in leaves where it is associated with the induction of numerous catabolic and proteolytic processes that participate in the degradation of macromolecules and ultimately in the dismantling of organelles, allowing the release of numerous macro and microelements.

As such, leaf senescence related metabolism is essential for the remobilization of nutrients from the leaves towards the growing organs and especially to the seeds during the reproductive phase (Gregersen et al., 2013; Avice and Etienne, 2014; Diaz-Mendoza et al., 2016). Leaf senescence is known to be regulated by numerous transcription factors belonging to the WRKY, NAC and ERFs families (Cao et al., 2023) that control the switch from anabolic to catabolic metabolisms and the expression of the senescence associated (SAGs) and senescence down regulated (SDGs) genes. Many cysteine related proteins and autophagy genes are represented in the SAGs lists (Avice and Etienne, 2014; Havé et al., 2017). SDGs are mainly represented by photosynthesis associated genes, protease inhibitors, and transcription factors related to chloroplast maintenance as the GLK1 proteins (Breeze et al., 2011).

Although roots can represent substantial nutrient reservoir in many plants as perennials, meadow plants, rhizome and taproot plants, root metabolism during senescence and the regulation of root senescence remain largely under-investigated. Recently, Wojciechowska et al., (2017) reported the induction of several N and amino acid transporters and of glutamate catabolic enzymes in the roots of *Populus trichocarpa* during senescence.

The few reports dealing with root aging describe root senescence as an age-dependant process leading to root browning, reduction in nutrient uptake from the soil, and programmed cell death in the root cortical cell layer (Eissenstat, 2000; Bingham, 2007; Schneider and Lynch, 2018). More recently, it was shown that barley roots undergo a genetically determined intrinsic senescence program, mainly influenced by plant age and sharing features with leaf senescence (Liu et al., 2019). Similarities with leaf senescence resides in the implication of the same transcription factor families NAC, WRKY, and ERF in the control of root senescence. The loss root physical integrity is likely modulated by abscisic acid and cytokinin like in leaves. James et al., 2018a showed that when Arabidopsis is facing nitrate limitation, the remobilization of root N is crucial for grain filling. This remobilization of N from Arabidopsis roots to the seeds is concomitant with root protein degradation and with the appearance of the SAG12 cysteine protease in root tissues (James et al., 2019). Rapeseed is a N demanding plant with mediocre capacity for N remobilization. It has the particularity of having a compartmentalized root system divided in a taproot and lateral roots. Several studies on rapeseed nutrient fluxes showed that taproot is an important reservoir of nutrient available for remobilisation during the monocarpic senescence (Rossato et al., 2002b; Gombert et al., 2010; Girondé et al., 2015). Rossato et al., (2002b) showed that nitrate uptake decreases in rapeseed roots during aging and that several proteins and especially a 19 KDa vegetative storage protein (VSP) located in the taproot of rapeseed are hydrolyzed in parallel of an increase of N export to the reproductive organs. The lateral roots are efficient at N uptake from the soil and their decay during reproductive stage results in the decrease of N uptake while at the same time, seed filling requires substantial N input. The taproot, which has an N-storage function, is then solicited for remobilization of the nutrients towards the new sinks. The aim of the present study was to investigate rapeseed root senescence and to identify the main proteases involved during the N recycling phase in comparison with what has been observed in the senescence of Arabidopsis leaves.

Rapeseed plants were grown in hydroponic and submitted or not to long-term nitrate starvation to modulate senescence severity during the reproductive stage. Whole plant phenotype and lateral root and taproot metabolism were investigated at three stages of the reproductive stage that correspond to the end of the vernalization period, the appearance of flowering buds, and the pod developping stage. N budgets, root structure and shotgun proteomics highlight the biological processes that are the most affected during root senescence in each root compartment.

## Material and methods

### Plant material

Seeds of *Brassica napus* var. Aviso were germinated on perlite over demineralized water for 4 days in the dark followed by 6 days under natural light. After the emergence of the first true leaf, seedlings were transferred to a 10 L tank (ten seedlings per tank) containing the following nutrient solution: 3.75 mM NO_3-_ nutrient solution (1.25 mM Ca(NO_3_)_2_.4H_2_O, 1.25 mM KNO_3_, 0.5 mM MgSO_4_, 0.25 mM KH_2_PO_4_, 0.2 mM EDTA.2NaFe.3H_2_O, 14 μM H_3_BO_3_, 5 μM MnSO_4_, 3 μM ZnSO_4_, 0.7 μM (NH_4_)_6_Mo_7_O_24_, 0.7 μM CuSO_4_, 0.1 μM CoCl_2_) renewed every week for 22 days. At 32 days after sowing (DAS), plants were transferred under two contrasting N conditions: High Nitrogen (HN; 3.75 mM N) and Low Nitrogen (LN; 4.2 μM N) conditions and continue to grow in greenhouse with a thermoperiod of 20/17 °C day/night and a photoperiod of 16 h with average photosynthetically active radiation of 350 µmol photons m^−2^.s^−1^ at canopy height (natural light supplemented with high-pressure sodium lamps Philips, MASTER Green Power T400W, Amsterdam, Netherlands). At 47 DAS, plantlets were subjected to a 60 d period of vernalization in a climatic chamber maintained at 4 °C with artificial light (220 µmol photons m^−2^.s^−1^) during the day (10 h day/14 h night). At 108 DAS, vernalization was stopped and plants were transferred to the same growth condition as before the vernalization but each plant was grown in an individual hydroponic pot of 4 L for 57 days until the end of the flowering of principal stem (at 164 DAS). At 108 DAS (vernalization output; C1 T0), 130 DAS (beginning of flowering; F1; T1), and 165 DAS (pods development; G4; T2) plants were harvested. A portion of root tissues was frozen in liquid nitrogen and stored at -80 °C for further biochemical analysis, the remaining was stored in an oven (60 °C, 4 days) to obtain dry weight for biomass determination.

### Elementary analysis

All dried sampled of taproot and lateral roots were ground to a fine powder using stainless steel beads in an oscillating grinder (Mixer Mill MM400, Retsch, Haan, Germany). As detailed previously by Maillard et al., (2016), the iron (Fe) concentration was quantified after acid digestion of dry weight samples (about 40 mg) with high-resolution inductively coupled plasma mass spectrometry (HR-ICP-MS, Element 2TM, Thermo Scientific) and using internal and external standards. Concerning the nitrogen (N) concentration, they were quantified on 2 mg of dried powder with an elemental analyzer (EA3000, EuroVector, Milan, Italy).

### Extraction and quantification of soluble and insoluble proteins

Soluble proteins in McIlvaine buffer (McIlvaine, 1921) were extracted from 200 mg of frozen fresh root tissue ground in a mortar with 300 µL citrate-phosphate buffer (20 mM citrate, 160 mM phosphate, pH 6.8 containing 50 mg of PVPP). After centrifugation at 20,000 g at 4 °C for 20 min, the supernatant was recovered and the proteins are assayed. The pellet is kept to extract the insoluble proteins in McIlvaine buffer. The pellet was resuspended in thiourea/urea buffer. After 1 h of incubation with shaking at room temperature, the extracts were centrifuged twice at 20,000 g at 4 °C for 10 min and the supernatant with insoluble proteins was collected. The concentrations of the soluble and insoluble protein extracts were determined in the supernatants by protein staining (Bradford, 1976) using bovine serum albumin (BSA) as a standard.

### Determination of proteolytic activities

Protease activities were determined on the soluble protein extracts and with using the Abcam Assay Kit (ab111750), which incorporates fluorescein isothiocyanate (FITC)-labelled casein as a general protease substrate. Protease activities were determined at pH 5.5 and 7.5. For this, 15 μg of soluble proteins were incubated in a 200 µl reaction volume containing 2 mM dithiothreitol (DTT) and sodium-acetate buffer (50 mM, pH 5.5) for CPs and APs or Tris-base buffer (50 mM; pH 7.5) for SPs. Protease class activities were obtained by pre-incubating sample with the addition of 50 µM of protease class-specific inhibitor in dimethyl sulfoxide (DMSO): E-64 for CPs, aprotinin for SPs and pepstatin A for APs. After 30 min of incubation at 37°C, the fluorescence of peptide fragments was measured at an excitation/emission (Ex/Em) wavelength of 485/530 nm.

### Shotgun proteomic analyses

Protein extraction was performed using Phenol Extraction as describe in Belouah et al., (2020). Protein pellets were then solubilized with 6 M Urea, 2 M Thiourea, 30 mM TRIS-HCL pH 8.5, 10 mM DTT and 0.1 % Rapigest (Waters). Protein content were estimate using 2D QuantKit (GE Healthcare) and adjusted to 2 µg/µl. 20 µg of protein were digested and desalted as describe in Belouah et al., (2020).

A total of 400ng of desalted peptide digest were injected on Thermo Qexactive Plus (thermo) coupled to a Eksigent nanoLC ultra 2D (see supplementary method S1 for detailed parameter). Peptide identification was performed using Xtandem (piledriver 2015.04.01.1) against Brassica Napus refseq genome (https://www.ncbi.nlm.nih.gov/protein/?term=txid3708[Organism:exp]) and in house protein contaminant database (55 Entries) as describe in Balliau et al., (2018). The detailed parameters are described in supplementary method S2.

Protein inference and quantification were performed using i2masschroq software (http://pappso.inrae.fr/bioinfo/i2masschroq/, Langella et al., 2017; Valot et al., 2011). Protein Evalue was set to 0.00001 with 2 to distinct peptides with a Evalue of 0,01 resulting to a fdr of 0.1 at psm level, 0.11 at peptide level and 0.015 at protein level. Quantification was performed as describe in Balliau et al., (2018). For XIC quantification Peptide were kept if it present at least in 90% of samples, and if the correlation for all peptide depending to a protein have a correlation upper to 0.5. When the peptides of a protein were not present or not reproducibly observed in one or more conditions, spectral counting (SC) was used in place of XICs analysis. For SC quantification, proteins were kept if they are observed by a minimum of 5 spectrum in one sample. The mass spectrometry proteomics data have been deposited to the ProteomeXchange Consortium via the PRIDE (Perez-Riverol et al., 2022) partner repository with the dataset identifier PXD050894.

### Proteomic data: statistics and data mining

Statistical analyses were performed using Perseus software (https://maxquant.net/perseus/). Global proteomic data were analyzed using ANOVA with developmental stage as variable factor. For quantification of proteins by the XICs and SC method, statistical analyses were carried out on log2 transformed protein abundance. Proteins with a student test FDR ≤0.05 were considered to be significantly differentially accumulated. Missing data for the proteins analyzed by SC method were replaced from normal distribution No FC threshold was applied because a small variation in protease abundance can have a large impact on the proteome due to post-translational regulation and the large spectrum of protease substrates. Proteases were identified using the MEROPS database (https://www.ebi.ac.uk/merops/) and Gene Ontology (GO).

The GO and predicted subcellular localization were provided by the panther database (pantherdb.org) and SUBA5 site (https://suba.live/), respectively. The heatmap representation was performed with proteases that are differentially abundant over time with a pearson clustering method using the ComplexHeatmap R package (v2.13.3). GO enrichment analyses with the *Brassica napus* L. genome as a background were performed with using Cytoscape (v3.9.1) plug-in ClueGO (v2.5.9 ; Fischer test with FDR correction) (Bindea et al., 2009) or Pantherdb (https://www.pantherdb.org/ ; Fischer test with FDR correction) (Mi et al., 2019). GO enrichment analyses with Arabidopsis genome as a background were performed with VirtualPlant1.3 (http://virtualplant.bio.nyu.edu/cgi-bin/vpweb/ ; Fischer test with FDR correction) (Katari et al., 2010) Results focused on terms identified with a p-value < 0.05. The ClueGO network and grouping were built using kappa statistics to reflect the relationship between terms according to the resemblance of related proteins. Functionally grouped networks with terms as nodes were connected based on their kappa score level (≥ 0.4).

### Determination of Amino Acid Content

Amino acids were extracted from 100 mg of dry matter to which 1 mL of 80 % ethanol was added. Samples were incubated under agitation at 80°C for 15 min, then centrifuged at 5,000 rpm at RT for 10 min. The supernatant was reserved. The pellet was resuspended in 1 mL of deionized water and incubated under agitation at 60 °c for 15 min followed by centrifugation at 5,000 rpm at RT for 10 min and the supernatant was collected with the previous one. This last step with 1 mL of deionized water was repeated a second time. The collected supernatant was dried in a concentrator. The pellet was re-suspended in 100 µl of water deionized. To 100 µl of extract, 1 mL of ninhydrin (2,2-dihydroxyindane-1,3-dione) reagent (3.9 mM SnCl_2_.H_2_O, 2 % ninhydrin (w/v), 100 mM citrate buffer pH 5.5 and 50 % DMSO (v/v)) was added and incubated at 100 °C for 20 min. After stopping the reaction with ice, the samples were diluted with 5 mL of 50 % ethanol and the absorbance at 570 nm was measured using glycine as a standard.

### Quantification of starch

Starch content was extracted from 50 mg of dry matter and analyzed using the “Total Starch” enzymatic kit (Megazyme International, County Wicklow, Ireland). Briefly, starch was digested first with thermal stable α-amylase and then with amyloglucosidase after gelatinization at 100 °C. Residual glucose was determined spectrophotometrically at 510 nm using glucose oxidase/peroxidase, and 4-aminoantipyrine (GOPOD reagent) and glucose as a standard. Weight of free glucose was converted to anhydroglucose using a multiplication factor of 162/180.

### Microscopy observation

The lateral root tissues were included in low melting point agarose (5%; w/v) and cut with a vibratome (Microm 650v; Thermo Scientific; United States) before observation with a light microscope (AX70 Olympus and Olympus SC30 camera, Japan) with the help of cellSens software.

The strong lignification of taproot tissues did not permit to use vibratome as for lateral roots. Consequently, sections of 2 mm of taproot were fixed with 2.5 % glutaraldehyde in phosphate buffer 0.1 M pH 7 from 1 hour to several days at 4 °C. The sections were rinsed in Phosphate buffer 0.1 M pH 7 three times and post-fixed 2 hour with 1 % osmium tetroxyde in 0.1 M phosphate buffer pH 7, the sections were rinsed in Phosphate buffer three times. The cells were then dehydrated in progressive bath of ethanol (70-100 %), embedded in resin EMbed 812 and polymerised 48 h at 60 °C.

For tissue observation, 1 µm semi-fine sections with ultramicrotome stained with 0.5 % toluidine blue (1 % sodium borate) were made to be observed with a classical light microscope (Olympus AX70, and Olympus SC30 camera, Japan).

For ultrastructure observation ultrathin sections of 80 nm were done and contrasted with uranyl acetate and lead citrate. The sections were observed with transmission electron microscope JEOL 1011 and images were taken with Camera gatan Orius 200 and a digital micrograph.

### Statistical analysis

For all parameters, at least three biological replicates were measured (n ≥ 3). All data are presented as mean ± standard error (SE). To compare different data between different times or treatments, Tukey tests were performed after verifying compliance of normality with Shapiro-wilk test with R software. When the data did not conform to the normal distribution, they were transformed into Log. Statistical significance was postulated at p ≤ 0.05.

## RESULTS

### Phenotype of plants subjected or not to a N deprivation

Irrespective of developmental stages (T0, T1 or T2), the LN plants developed less leaves, with smaller size and reddish phenotype compared to the HN plants (Figure 1). At flowering stage (T1), the number of flowers was lower on LN than on HN plants. Still, all the HN and LN plants produced pods (Figure 1). Both lateral roots (LR) and taproot (T) exhibited increasing brownish colour over time, especially striking at the pod filling T2 (Figure 1).

**Figure 1.**
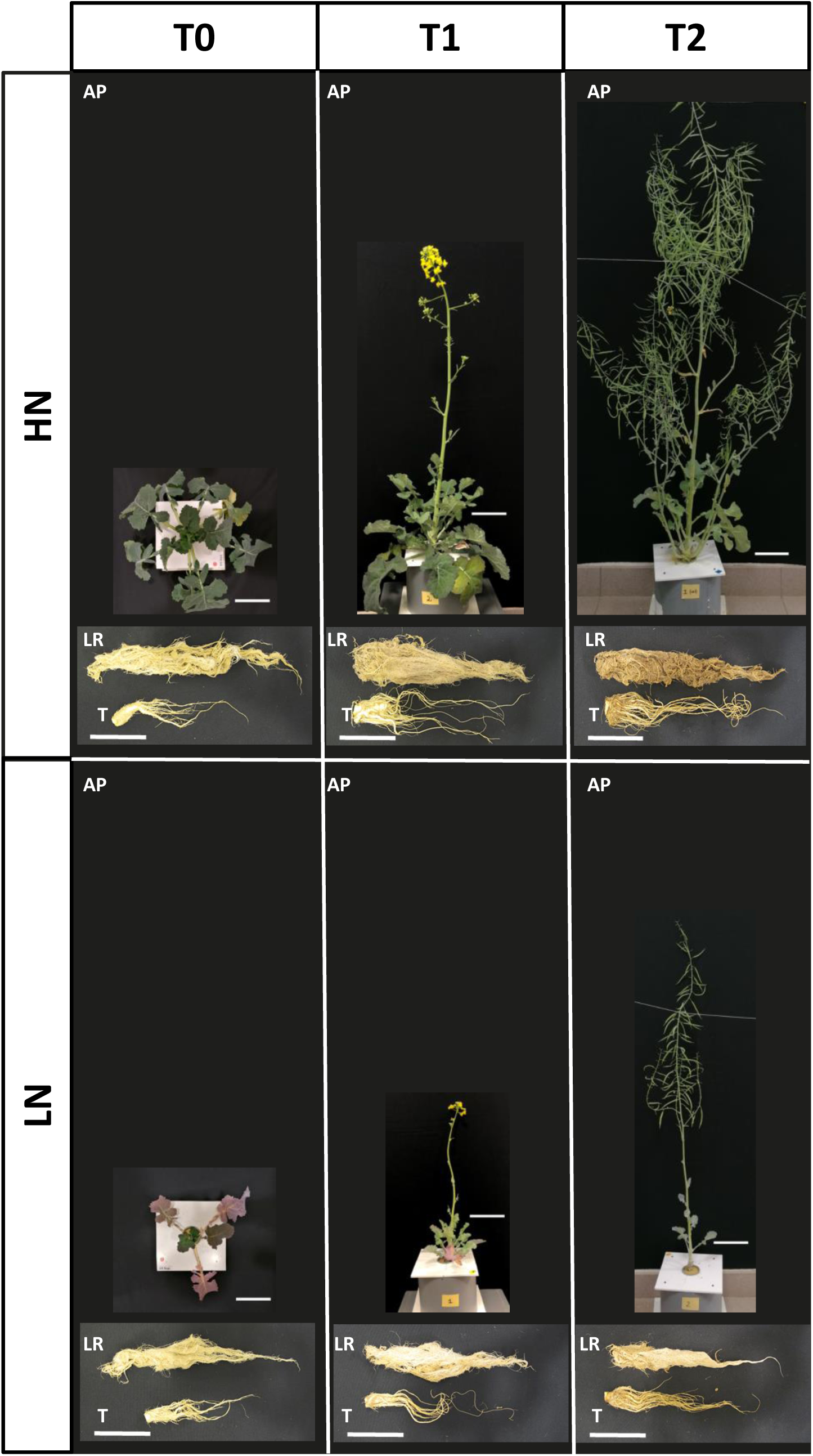
Development of rapeseed cultivated under high (HN) and low nitrogen (LN) conditions. The different harvest times T0, T1 and T2 correspond to vernalization output (B9), beginning of flowering (F1) and pod developments (G4) stages, respectively. AP: Aerial Parts, LR: Lateral Roots, T: Taproot. The scale bars represent 10 cm.

The smaller size of LN plants observed in Figure 1 was in good accordance with the 5 fold lower biomass of LN plants compared to HN plants at T2 (17.58 vs 95 g; Figures 2A and 2B) and was the result of the limitation of the biomass of all the organs by nitrate starvation, with the exception of lateral root biomass at the vernalization exit stage (T0), which is similar regardless of the N treatment (Figures 2A and 2B).

**Figure 2.**
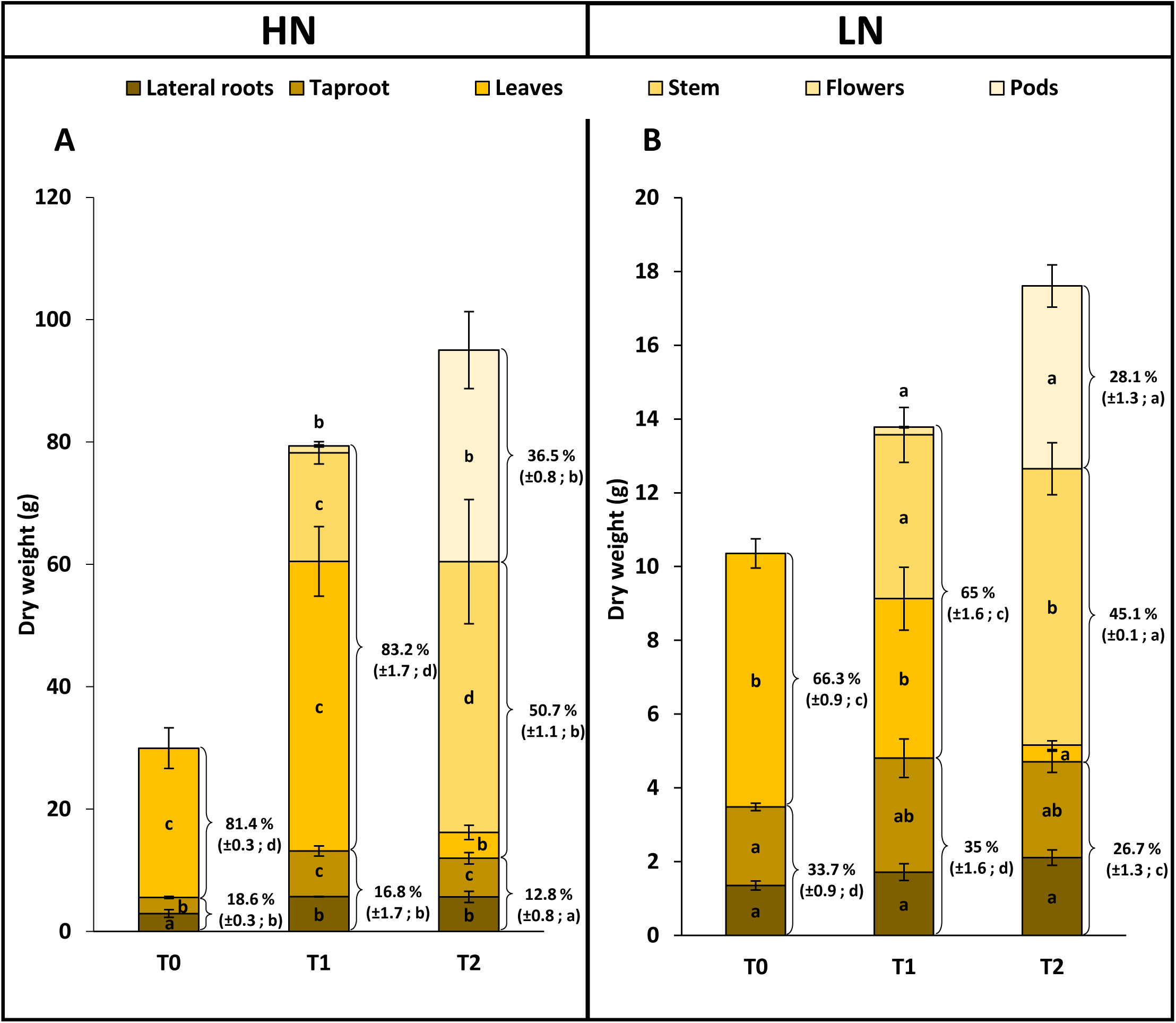
Biomasses of whole plant and of the different compartments of rapeseeds cultivated under high (HN; A) or low nitrogen (LN; B) conditions. The different harvest times T0, T1 and T2 correspond to vernalization output (B9), beginning of flowering (F1) and pod developments (G4) stages, respectively. Each colour in a histogram bar correspond to the biomass of each plant compartment (means ± SE; n=3). The brackets to the right of the histogram bars correspond to the root (lateral roots and taproot), aerial (leaves, stems, flowers) and pods parts and the associated value are the percentage (± SE; n=3) of each part relative to the total biomass. For a given compartment or a given part of the plant and regardless to the nitrogen condition, different letters indicate significant differences between developmental stages (p-value ≤ 0.05).

Although plant and organ biomasses were significantly lower under LN than under HN, the partitioning of biomasses in root was twice higher in LN plants than in HN plants (33.7 vs 18.6 at T0; 35 vs 16.8 at T1 and 26.7% vs 12.8 % at T2 respectively; Figures 2A and 2B). This shows that under N starvation shoot biomass was neglected compared to root biomass.

### Anatomical analysis of taproot and lateral roots

Lateral root tissues were observed under light microscope after Evan blue staining to monitor cell viability (Figure 3). For a given stage (T0, T1 or T2), no anatomical differences between the roots of LN and HN plants can be observe. At T0, which corresponds to the exit of vernalization, root section was a nice circle and the LN and HN roots mainly differed by root diameter, the size of the cells and the number of cell layers in the cortex (4-5 for HN, 3 for LN). There was no evidence of cell death (Figures 3A and 3D). At the beginning of flowering (T1) and pod-filling (T2) stages, the shape of cortical and epidermis cells was strongly affected, roundness was lost, especially in the outer layers and in HN root. The slight blue color of these cells presents at T1 and that amplifies at T2 suggests a loss of cell viability with aging (Figures 3B, 3C, 3E and 3F). It can be noticed that the integrity of the stele and vascular tissues seems by contrast well preserved during aging in both HN and LN roots.

**Figure 3.**
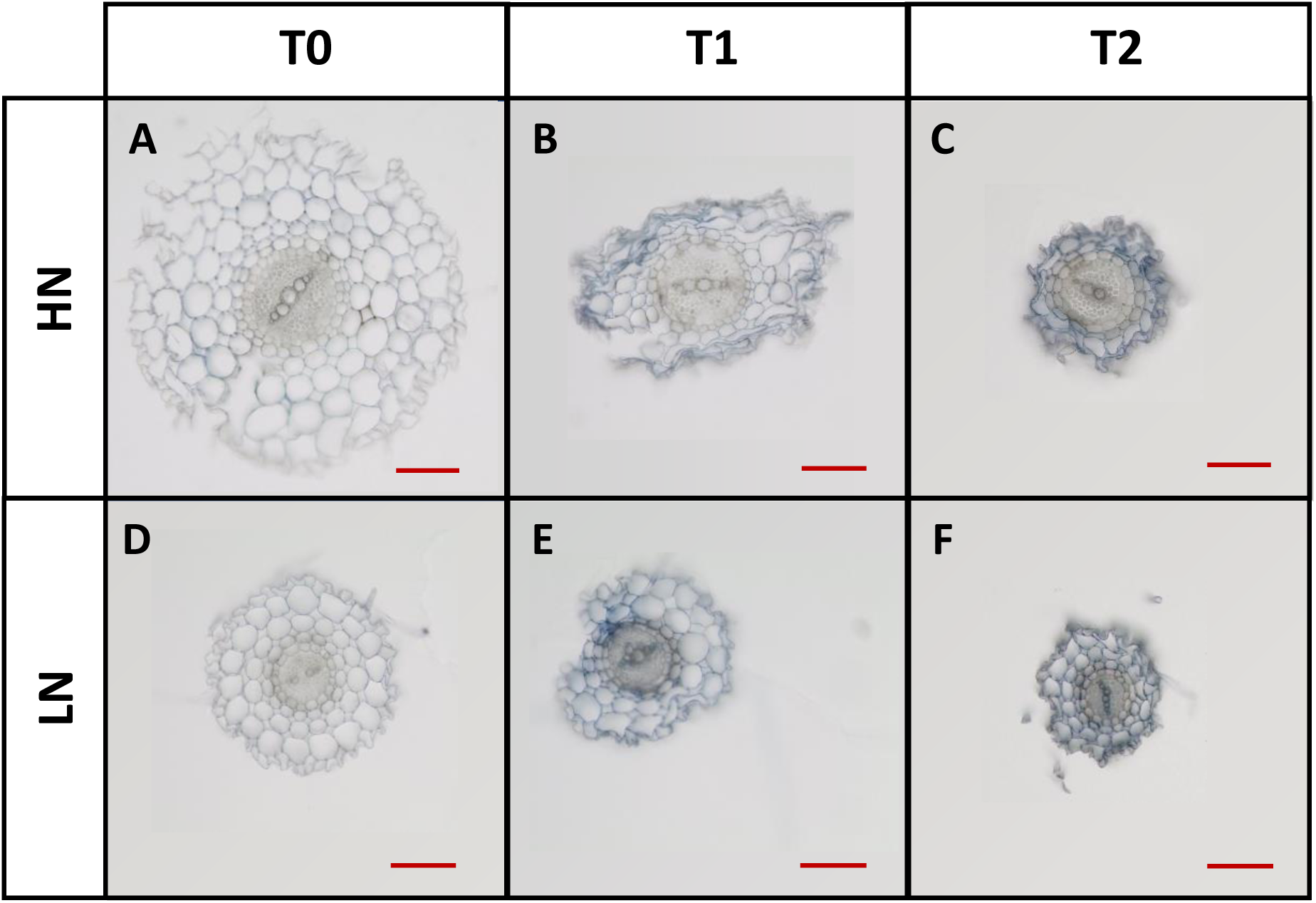
Structural changes in the cortex of rapeseed lateral roots over time under high (HN) or low nitrogen (LN) conditions. Pictures of root tissue sections, observed after staining with Evans blue using a light microscope, were obtained at vernalization output (T0; A, D), beginning of flowering (T1; B, E) and pod developments (T2; C, F) under HN (A, B, C) or LN (D, E, F) conditions. The red scale bars represent 100 µm. Representative pictures are shown n=3.

Observation of taproot tissues with light microscopy shows no change in the roundness of cell shape over time, but a decrease in cell cohesion in both HN and LN taproots (Figures 4A, 4B, 4C, 4G, 4H and 4I). Both LN and HN taproot cells contain numerous organelles, likely amyloplasts, whose numbers decrease between T0 (Figures 4A and 4G) and T1 (Figures 4B, 4H). These structures are completely disappeared at the T2 pod-filling stage (Figures 4C and 4I). Ultrastructure images of the LN and HN taproot cells confirm that these structures are amyloplasts that are intact at T0 with a high starch concentration in the stroma (Figures 4D, 4J) and undergo decay at T1 and T2. At T1, they have lost their structural integrity and starch concentrations, their size is strongly reduced and they present crystal structures that could not be observed at T0. At T2, amyloplasts are dramatically shrunken and appear as dislocated bodies that still contain numerous crystal structures but no more starch (Figures 4E, 4K, 4F and 4L). The crystal structures accumulating in degenerating amyloplast strongly resemble to the crystalloid ferritin structures previously observed in chloroplast of *Mesembryanthemum crystallinum* subjected to salt stress (Paramonova et al., 2007). They accumulate under both LN and HN conditions with aging (Figures 4E, 4K, 4F and 4L). This observation is in agreement with the increase in the iron concentration between T1 and T2 in both the HN and LN taproots (Figure S1).

**Figure 4.**
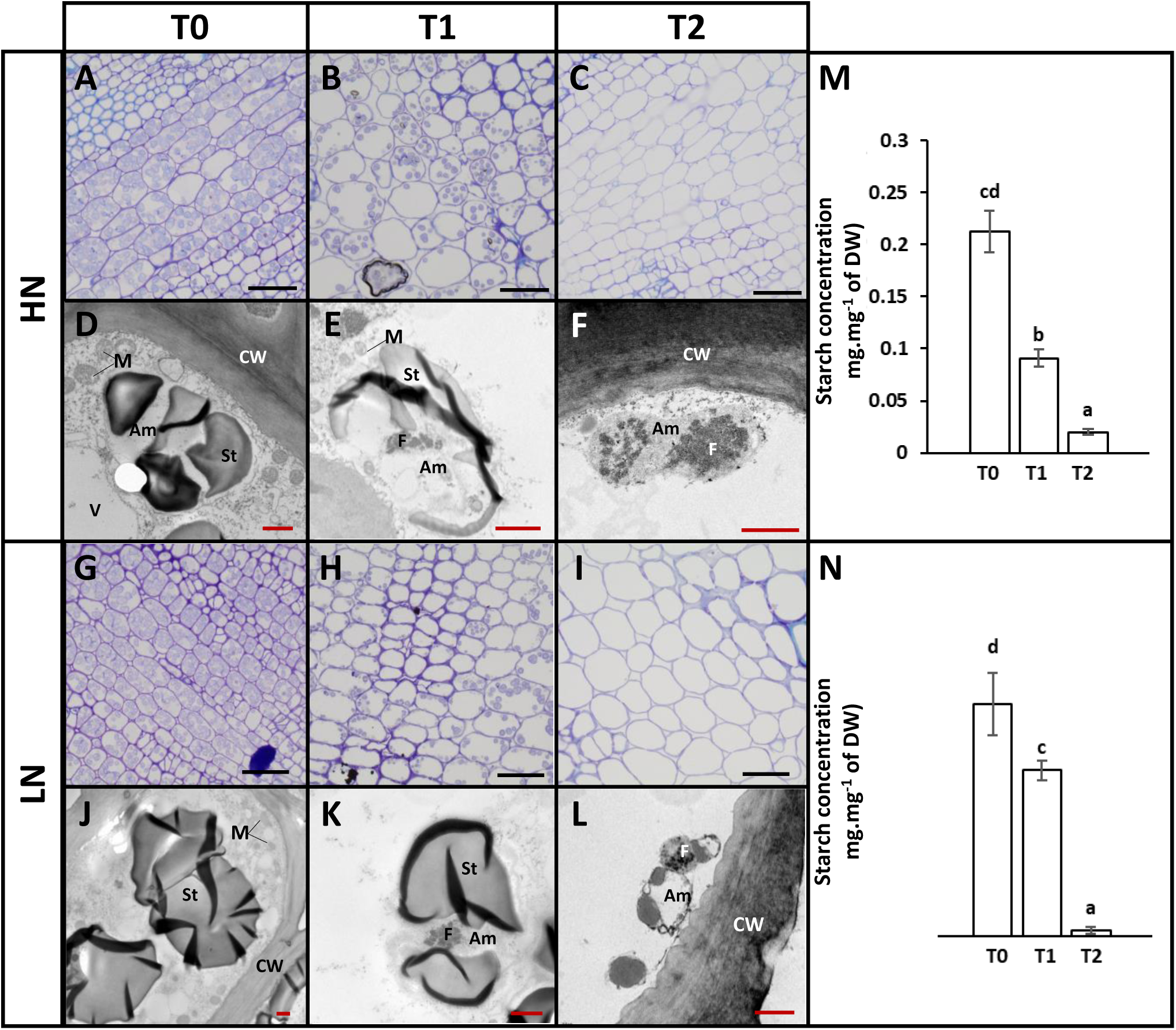
Structure, ultra-structure and starch concentration changes in the taproot tissues during the development of rapeseed cultivated under high (HN) or low nitrogen (LN) conditions. Taproot structure were obtained after staining with 0.5% toluidine blue (1% sodium borate) using a light microscope at vernalization output (T0; A, G), beginning of flowering (T1; B, H) and pod developments (T2; C, I) under HN (A, B, C) or LN (G, H, I) conditions. Cellular ultrastructure was obtained after staining with 1% osmium tetroxide by using a transmission electron microscope at vernalization output (T0; D, J), beginning of flowering (T1; E, K) and pod developments (T2; F, L) under HN (D, E, F) or LN (J, K, L) conditions. Black and red scale bars correspond to 50 µm and 1 µm, respectively. Am: Amyloplast, CW: Cell-wall F: Ferritin deposits, M: Mitochondria, St: Starch grain, V: Vacuole. Starch concentrations in taproot were measured over time under HN (M) or LN (N) conditions. Values are means ± SE (n=3). Significant differences between development stages regardless the N condition are indicated by different letters (p-value ≤ 0.05).

The decrease of starch concentration along amyloplast decay was confirmed by starch measurement assays (Figure 4M and 4N). At vernalization exit stage (T0), start concentrations were similar in HN and LN taproots (21.3 % and 22.3 % of total taproot biomass, respectively). Start concentrations decreased with aging in both LN and HN taproots but more sharply in the HN tissues, then representing at T2 only 2 % and 0.5 % of the taproot dry biomass obtained under the HN and LN conditions, respectively (Figures 4M and 4N).

### Evolution of nitrogen, proteins and amino acid contents in taproot and lateral roots

The amount of total N, proteins (soluble and insoluble) and free amino acids were higher in the lateral roots and taproots of HN plants than in LN plants at all the harvesting times (Figures 5A-5F).

**Figure 5.**
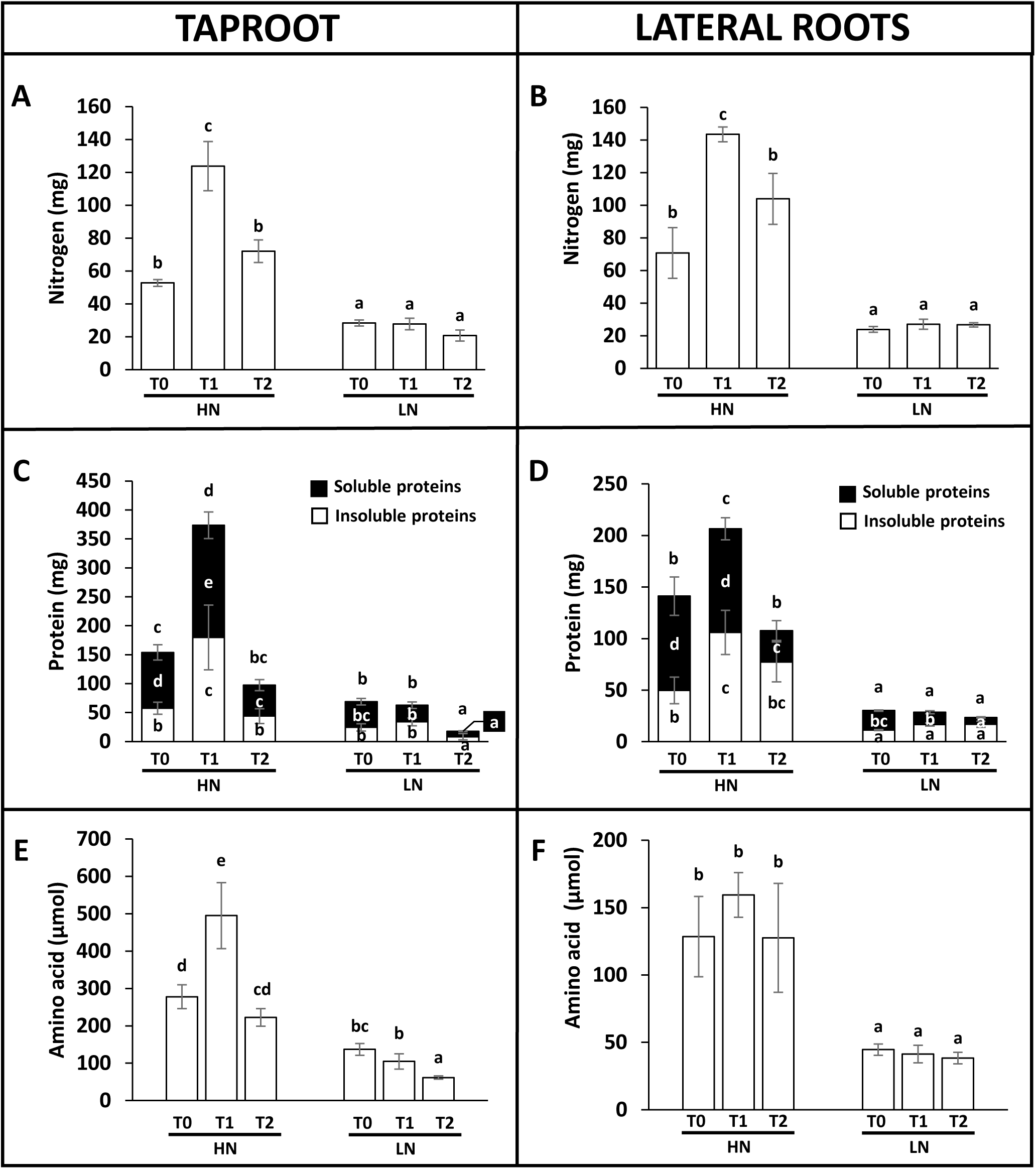
Total nitrogen, protein and amino acid amounts in the taproot and lateral root tissues during the development of rapeseed cultivated under high (HN) or low nitrogen (LN) conditions. The different harvest times T0, T1 and T2 correspond to vernalization output (B9), beginning of flowering (F1) and pod developments (G4) stages, respectively. Total nitrogen (A, B), soluble and insoluble proteins (C, D) and amino acid (E, F) amounts were measured in taproot (A, C, E) and lateral roots (B, D, F) over time under high (HN) or low nitrogen (LN) conditions. Values are means ± SE (n = 3). For a given root tissue, different letters indicate significant differences (p-value ≤ 0.05) regardless to the N condition and developmental stages.

In the lateral roots and taproots of HN plants, the amount of total N increased 2-fold between T0 and T1 (Figures 5A and 5B), and was reduced by one third for taproot and one fourth for lateral roots between T1 and T2. In contrast, in LN plants, the amount of N was globally maintained constant (c.a. 25 mg per organ) all along aging in both taproot and lateral roots (Figures 5A and 5B).

At T0, total, soluble and insoluble protein amounts were approximately at the same level in the taproot and lateral roots of HN plants (c.a. 150, 95 and 55 mg respectively; Figures 5C and 5D). In the taproot of HN plants, the amount of soluble proteins increased by 2-fold and insoluble proteins by 3-fold between T0 and T1. Then soluble and insoluble proteins decreased by 70-75 % between T1 and T2 (Figure 5C). In lateral roots of HN plants, only insoluble proteins increased significantly between T0 and T1 (2-fold) compared to taproot (Figure 5C). In HN plants, the decrease in total proteins in lateral roots (by 48 %; Figure 5D) between T1 and T2 was lower than that of taproot (by 74%; Figure 5C). In LN plants, total protein amounts in lateral roots were maintained at the same level all along aging whereas in the taproot, the amount decreases between T1 and T2 (Figures 5C and 5D). Irrespective of the N treatment, it can be noticed that the decrease in total protein amount in lateral roots is mainly due to a decrease in soluble protein while in taproot both insoluble and soluble proteins amounts are equally decreased (Figures 5C and 5D).

Changes in free amino acid contents in the taproot (Figures 5E) are quite similar to that of proteins (Figure 5C). In taproot of HN plants, amino acids increased by 1.8-fold between T0 and T1 and then decreases by 2.2-fold between T1 and T2. Amino acids contents steadily decreased by 2-fold from T0 to T2 in the LN taproots, while it was maintained to the same level in the LN lateral roots (Figure 5F). There was no significant change for amino acid contents in HN lateral roots along aging (Figure 5F).

### Proteases activities in taproot and lateral roots

Total and specific activities of different protease classes (cysteine, aspartic and serine) were measured at pH 5.5 or 7.5 in taproots and lateral roots (Figures 6). Total and specific proteases activities at pH 5.5 and 7.5 were always lower in the taproot than in lateral roots (Figures 6A and 6F). For taproot, all the activities measured at different pH and for total or specific protease families were not significantly different between HN and LN conditions, irrespective of plant age (Figures 6A-6E). All these activities show large increase at T2 while they were very low at T0 and T1 (Figures 6A-6E). Activity profiles were not the same in lateral roots as in taproot since total and specific protease activities increased progressively with ageing (Figure 6F-6J). However, for lateral roots, activity profiles are different in LN and HN plants. Irrespective of pH, protease classes and N treatment, activities are significantly induced between T0 and T1 in HN plants. However, the strongest protease activity change observed in lateral roots under LN is a remarkable increase between T1 and T2 (Figures 6F and 6I). In taproot and lateral roots, it can be noticed that total protease activites measured at the two pH, are mainly the results of the cysteine and aspartic protease activities at pH 5.5 and the serine protease activities at pH 7.5 (Figure 6).

**Figure 6.**
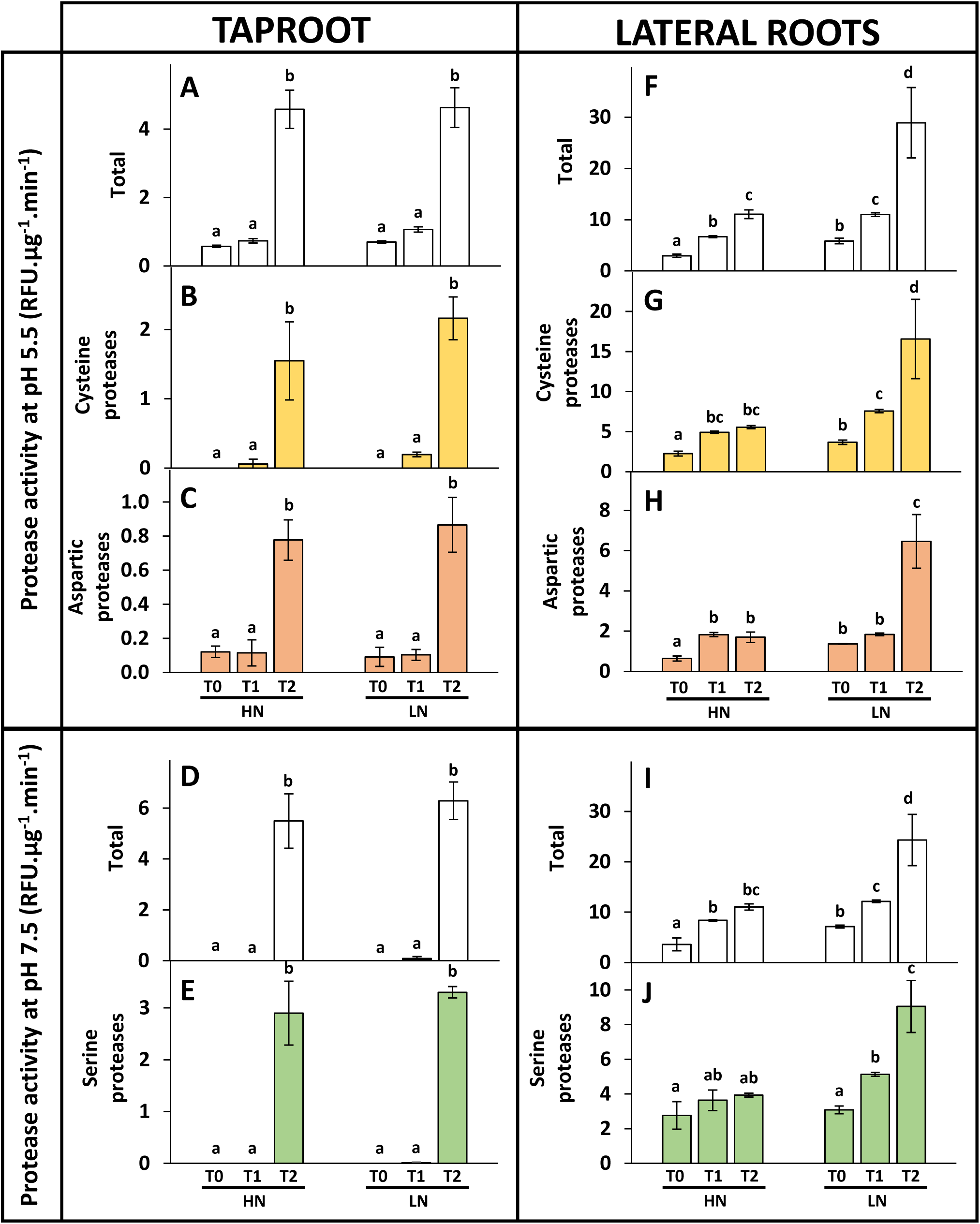
Total and specific proteases activities in the taproot and lateral root tissues during the development of rapeseed cultivated under high (HN) or low nitrogen (LN) conditions. The different harvest times T0, T1 and T2 correspond to vernalization output (B9), beginning of flowering (F1) and pod developments (G4) stages, respectively. The total proteases activities at pH 5.5 (A, F) and pH 7.5 (D, I), the cysteine (B, G) and the aspartic (C, H) protease activities at pH 5.5 and the serine protease activities at pH7.5 (E, J), were measured in taproot (A-E) and lateral roots (F-J) over time under HN and LN conditions. Protease activities are expressed in relative fluorescence units (RFU) per µg of protein per minute. Value are means ± SE (n=3). For a given root tissue, different letters indicate significant differences (p-value ≤ 0.05) regardless to the N condition and developmental stages.

### Proteomic analysis on taproot and lateral roots of plants grown under LN condition

In order to identify root senescence markers and provide a comprehensive picture or root ageing, shotgun proteomic analyses were performed on the taproot and lateral roots of plants grown under LN at the three time points previously analyzed (Table S1-S4). We focused on LN as this condition displayed the highest changes in protease activities and should provide a good picture of senescence and nutrient remobilizing processes (Figure 6).

The shotgun proteomic identified a total of 3028 and 2523 (Table S5 and S6) significantly differentially accumulated proteins (DAPs) depending on developmental stages in the taproot and lateral roots, respectively (ANOVA; FDR ≤0.05). Investigation of gene ontology using Cytoscape (v3.9.1) and the plug-in ClueGO (v2.5.9) (Bindea et al., 2009) identified 29 significant biological processes for the taproot and lateral root DAPs, the most significant of which are presented in Figures 7A and 7B. Twelve and thirteen over the twenty-nine biological processes identified are tightly related to N metabolism for the taproot and lateral roots, respectively. Among them, height are common to both root organs, four are specific to taproot and five to lateral roots. Interestingly, four enriched biological processes related to catabolism and organonitrogen compounds metabolism include proteases, which some were identified in the senescence of *B. napus* leaves by Poret et al., (2019) (Tables S7 and S8). Detoxification processes, which are consistent with senescence and catabolism are specifically enriched in the taproot and are also found in lateral roots but outside the 29 most enriched. (Figure 7A and table S8). As such taproot and lateral roots share several biological processes related to organic acid metabolic process (oxoacid and carboxylic acid metabolic processes), amino acids biosynthesis, nucleobase biosynthesis and catabolism.

**Figure 7.**
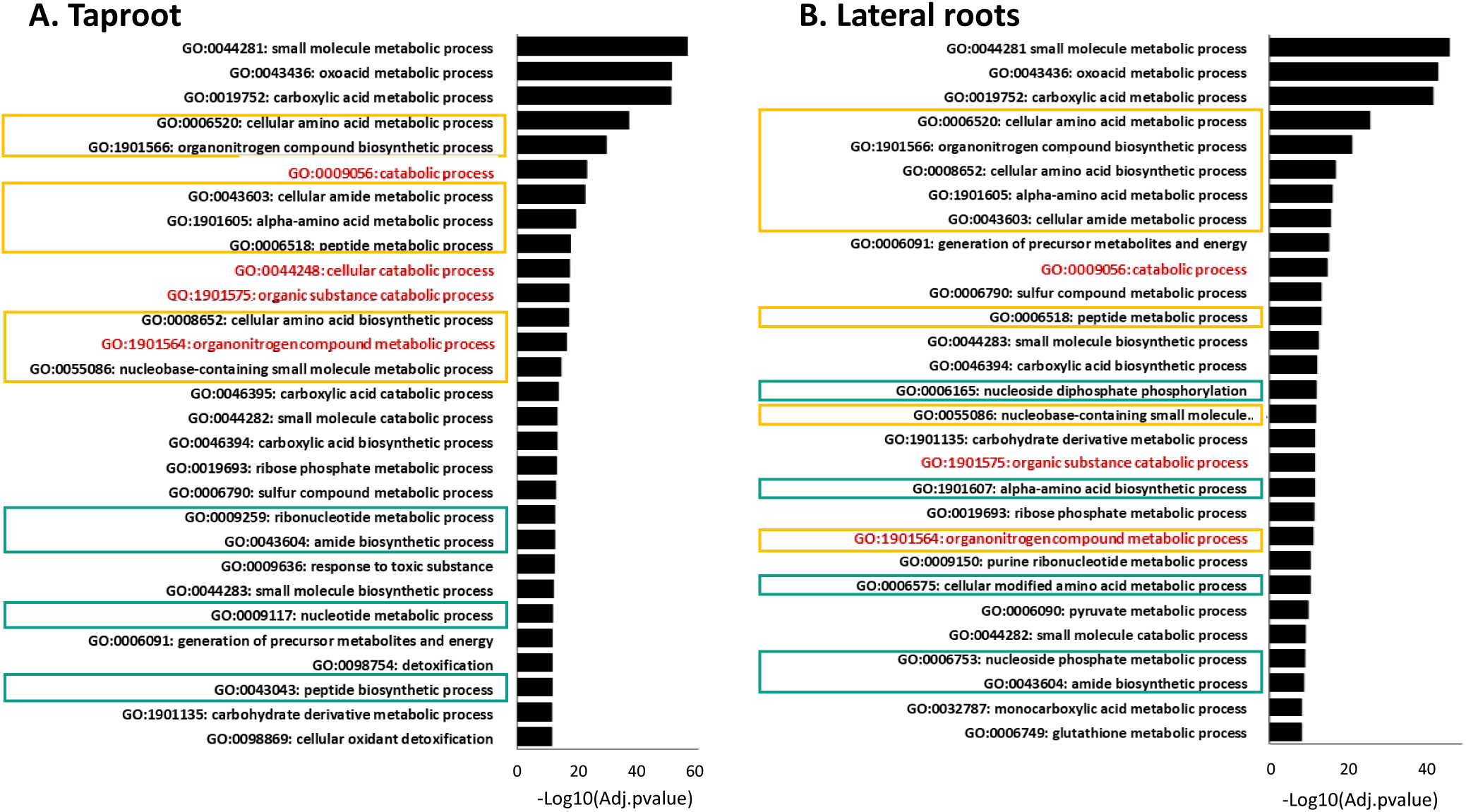
Biological processes in the taproot (A) and lateral roots (B) tissues during the development of rapeseed cultivated under low nitrogen (LN) conditions. Analysis of biological processes categories significantly using Cytoscape (v3.9.1) and the plug-in ClueGO (v2.5.9) (Bindea et al., 2009) enriched for the taproot (A) and lateral root DAPs was performed and the 29 most significant GO terms are presented. Categories related to N metabolism common or specific to root organs are framed in yellow and green, respectively. Enriched biological processes written in red include many proteases known to be involved in leaf senescence.

In order to get better insight of proteomic changes in taproot and lateral root, clustering was performed. Heatmaps identified four main groups for both taproot and lateral roots that share similar patterns (Figures 8A-8C; Table S5 and S6). The Clusters 1 correspond to proteins whose abundance decreases lately at T2 (T1≥T0>T2). The Clusters 2 correspond to proteins whose abundance decreases early since T1 (T0>T1>T2). The Cluster 3 include proteins with quite similar abundance at T0 and T1 and whose abundance significantly increase at T2 (T0-T1<T2). The Cluster 4 correspond to proteins whose abundance increase early at T1 and continue increasing at T2 (T0<T1<T2). Thus clusters 1 and 2 correspond to protein significantly depleted with senescence (SDP; senescence depleted protein) and clusters 4 and 5 of proteins significantly accumulated with senescence (SAP; senescence associated protein). Proportions of SDP and SAP are almost similar irrespective of root organs (Figures 8A and 8B).

**Figure 8.**
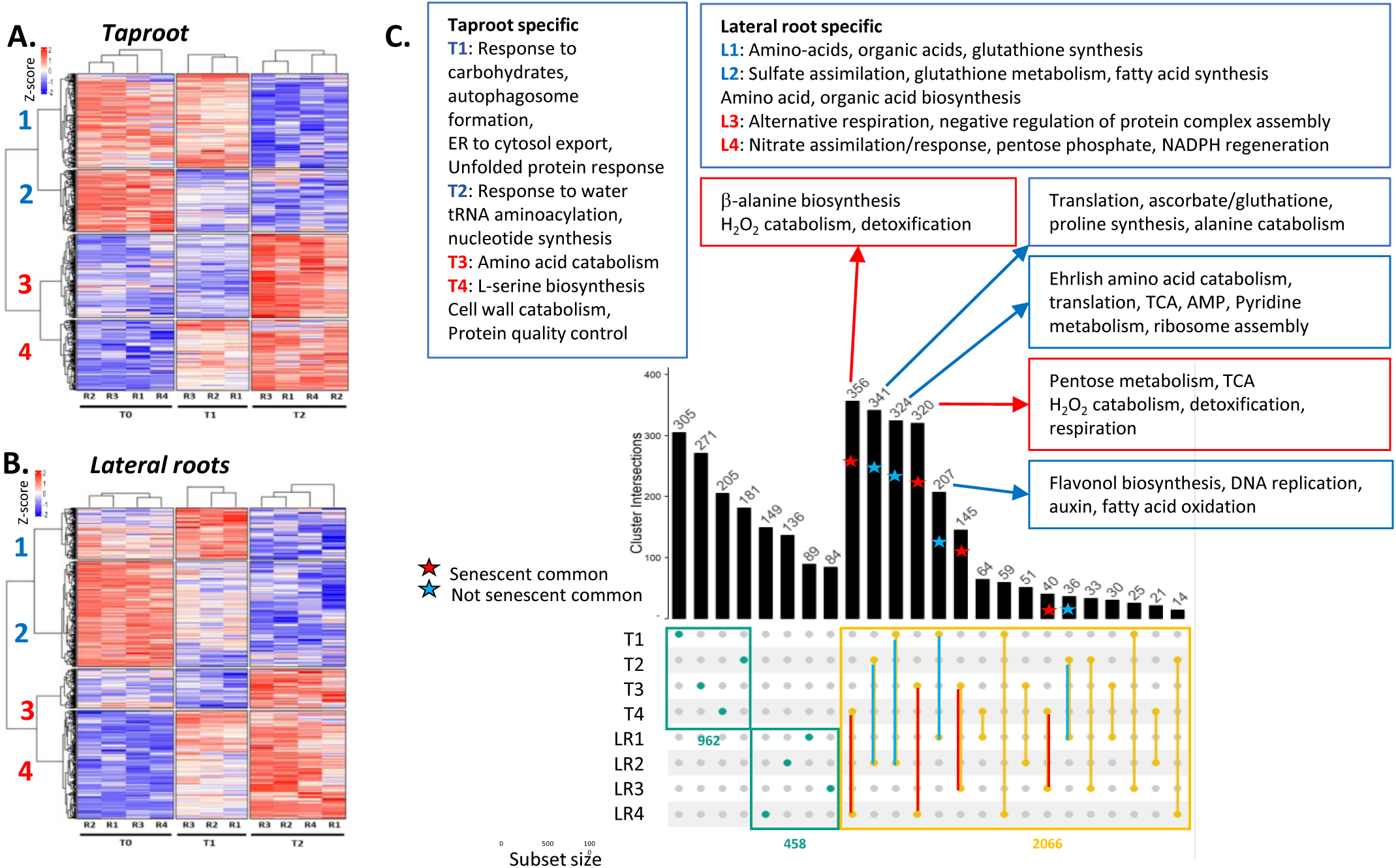
Changes in biological processes with taproot and lateral roots ageing. The different harvest times T0, T1 and T2 correspond to vernalization output (C1), beginning of flowering (F1) and pod developments (G4) stages, respectively. Heatmaps (A, B) represent the abundance of proteins (transformed in Z-score; red and blue for high and low accumulation, respectively) whose variability over time was confirmed by ANOVA in taproot (A) and in lateral roots (B). Each column corresponds to a biological replicate and the numbers at the left of the heatmap correspond to a cluster number. (C) UpSet plot identifies the time and organ specific DAPs (green spots) and those shared by taproot and lateral root (yellow spots). Senescence-related and early-development (not senescent) common DAPs are identified by red and blue colors respectively. Biological processes enrichment was performed with all proteins presented in each UpSet plot group using Panther (https://www.pantherdb.org/). Significant enrichment (Fischer test with FDR correction) are presented. According to (A) and (B) subsets used for Upset plots in (C) are: T1, taproot cluster 1; T2, taproot cluster 2; T3, taproot cluster 3; T4, taproot cluster 4; L1, lateral roots cluster 1; L2, lateral roots cluster 2; L3, lateral roots cluster 3; L4, lateral roots cluster 4.

UpSet plot representation of the taproot and lateral root clusters distinguish the proteins that are present in only one cluster and thus are organ and time specific, from the proteins that are found in two different clusters so in two different root organs (Figure 8C). The majority of the proteins present in two different clusters are senescence-depleted proteins (SDP) or senescence-associated proteins (SAP) common to the taproot and lateral root tissues. Only few are SDP in taproot and SAP in lateral root, or vice versa. Biological processes enriched in the different situations presented in figure 8C, and listed in supplemental dataset S9, reflect that the catabolism of amino acid, cell wall and peroxide and detoxification were increased with aging, while anabolic processes, translation, DNA replication were more active in young roots. We can notice that terms related to inorganic nitrate and sulfate assimilations were over-represented in the lateral roots which are the site of mineral uptakes.

### Identification of common root and leaf senescence markers

Proteomic data on leaf senescence would have been ideal when attempting to compare root senescence and leaf senescence. However, leaf senescence has been mainly studied using transcriptome analyses, and we could not find any large proteomic dataset from senescing leaves in public libraries. We then decided to explore the transcriptomic datasets and choose the transcriptome from Breeze et al. (2011) on Arabidopsis to determine whether root and leaves could share similar senescence markers. The transcriptome analysis of Breeze et al., (2011) provides gene expression at 11 time points along leaf development, and the spline analysis performed by authors on gene expressions permitted rigorous classification of senescence associated genes and senescence repressed genes in different clusters. Using both gene expression and clustering from Breeze’s study, we could identify in the taproot and lateral root many SDPs and SAPs that were homologous to the Arabidopsis senescence-repressed genes (SRG) and senescence-associated genes (SAG) (Tables S5 and S6). For both organs, we found root SAPs corresponding to leaf SAGs, and root SDPs corresponding to leaf SRGs that represent senescent and non-senescent markers of both root and leaf, respectively (figure 9). We also found numerous homologous corresponding to SAPs in the roots of *B. napus* and to SRGs in Arabidopsis leaves that we supposed they are long life proteins (accepting the hypothesis that their genes are regulated in the same way in Arabidopsis and in *B. napus*). At the opposite we also found numerous *B. napus* root SDPs corresponding to Arabidopsis SAGs and we supposed they could correspond to short life proteins with higher turn-over in senescing tissues. Analysis of the terms associated to the different categories was performed using VirtualPlant1.3 (with FDR correction). In the SAPs-SAGs class that represents senescence markers common to leaf and root, nucleobase catabolism (PYD homologues involved in uracil, thymidine and pyrimidine degradation) and oxidoreduction are over-represented in both lateral roots and taproot. Cysteine proteases are overrepresented in the taproot SAPs-SAGs class (Figure 9). Longevity markers represented by the SDP-SRG class are enriched in proteins related to carbon metabolism. For the two other classes (SDPs-SAGs and SAPs-SRGs) similar enriched categories were found for taproot and lateral roots. The SAPs-SRGs class is enriched in proteins related to fatty acid catabolism and oxidoreduction. The SAGs-SDPs class is enriched in hydrolase functions in both taproot and lateral roots, in amino-acid and organic acid metabolism in taproot and organic acid transport in lateral roots. The most striking result is that proteins/genes related to nucleobase degradation represent the common feature of leaf and root senescence in this study.

**Figure 9.**
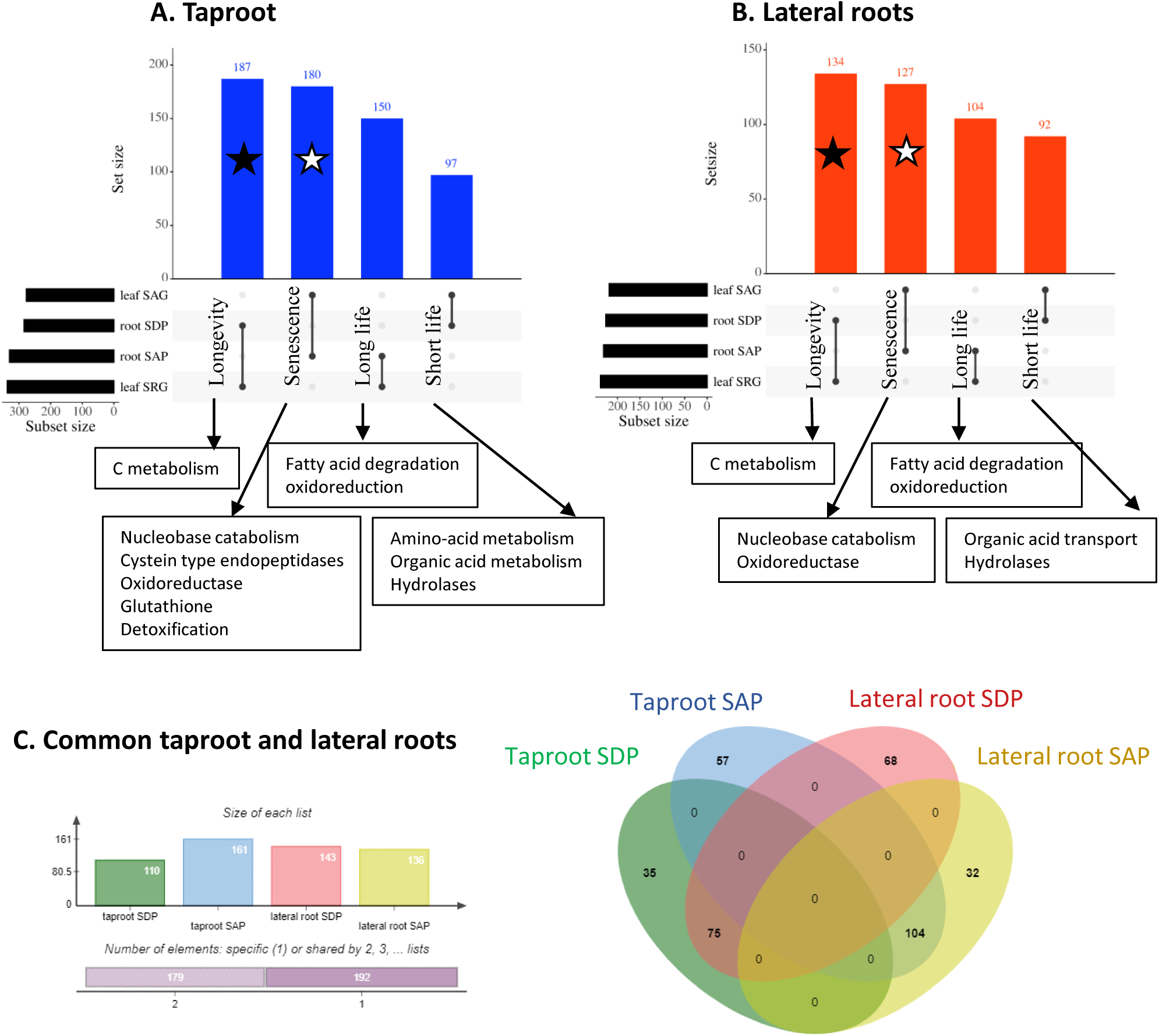
Longevity and senescence markers in taproot and lateral roots. Homologous genes of the *B. napus* senescence associated (SAP) and senescence depleted (SDP) proteins identified in taproot (A) and lateral roots (B) were identified amongst the senescence associated (SAG) and senescence repressed (SRG) genes from Breeze at al. (2023). The number of homologous genes corresponding to each category is indicated on top of each bar. The stars represent the common senescence markers (white stars) and the longevity markers (black stars) in *B. napus* roots and Arabidopsis leaves. Functional categories overrepresented in the set accessions are indicated next to the arrows (VirtualPlant1.3; FDR correction). (C) Venn diagram of the common markers (SDP and SAP) in taproot and lateral roots.

### Identification of proteases in taproot and lateral roots of plants grown under LN condition

Both protein decrease (Figure 5) and the increase in protease activities (Figure 6) prompted us to analyze in more detail the SAPs and SDPs related to the proteolytic activities (proteases and proteasome related proteins). A total of 165 and 112 proteases were identified in taproot and lateral roots respectively using the MEROPS database (https://www.ebi.ac.uk/merops/) and protease list was hand curated using gene ontology (Tables S10 and S11). According to the heatmaps of the taproot and lateral root proteases (Figure S2) we can globally define two senescence-accumulated (95 for taproot and 46 for lateral roots) and senescence-depleted (70 for taproot and 66 for lateral roots) protease groups (Figure S2 and Table S8-S9). UpSet plot presented in Figure 10 distinguish taproot specific proteases, lateral root specific proteases and proteases common to lateral roots and taproot. Very few common proteases were affected by aging in a different way in taproot and lateral roots, and we will not comment on them. Most of the common proteases were either over-accumulated or depleted with senescence in both taproot and lateral roots. The forty common senescence over-accumulated proteases were mainly represented by cysteine protease (13) and equal numbers of aspartate, serine and metallo-proteases. The forty common senescence depleted proteases were mainly represented by the proteasome subunits (16), metallo-proteases (12) and serine proteases. Such patterns were in good accordance with the senescence picture described for leaf senescence (Roberts et al., 2012).

**Figure 10.**
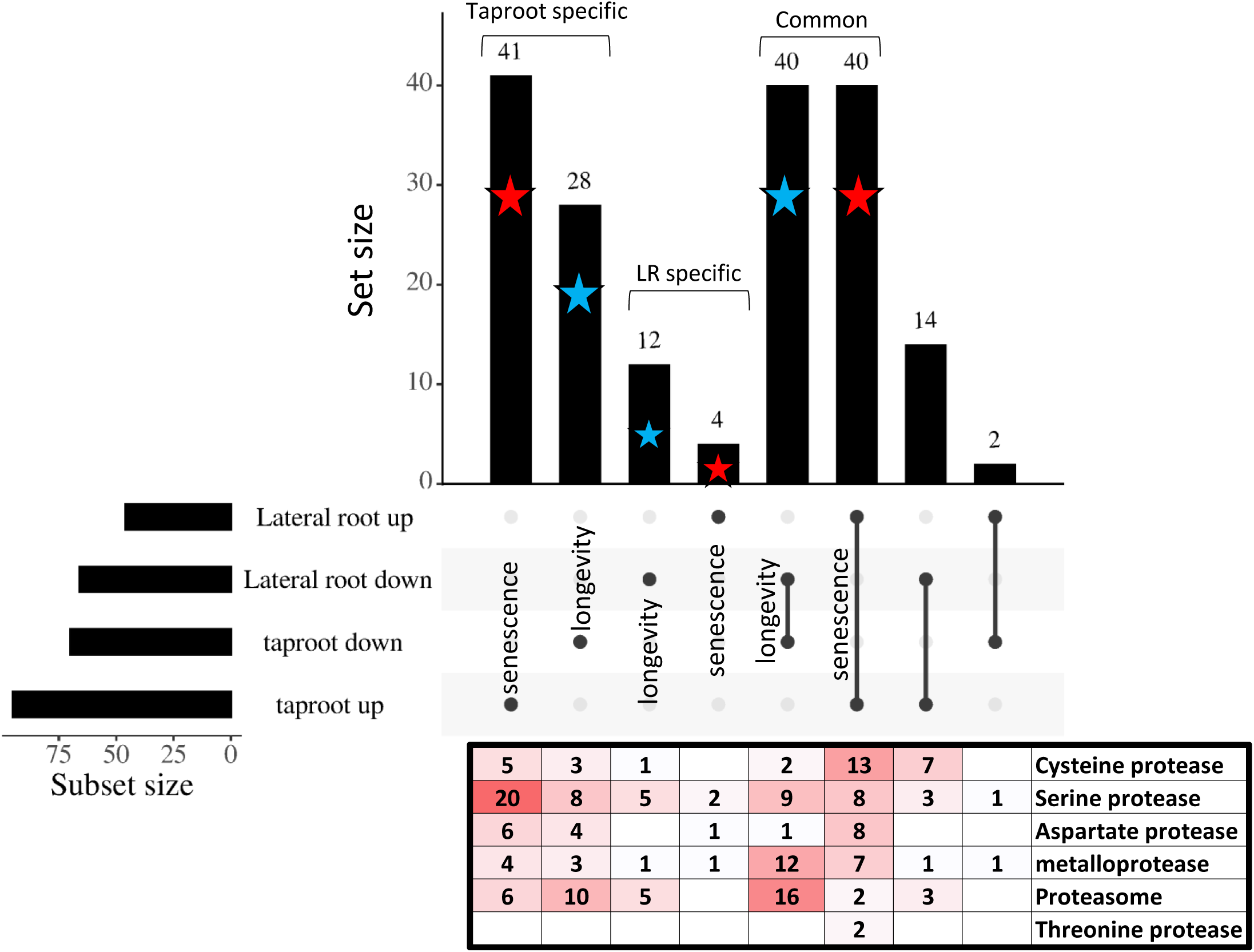
Senescence and longevity associated proteases in taproot and lateral roots. Proteases of taproot and lateral roots were qualified as senescence associated (up) or longevity associated (down) depending on if they were up-accumulated or depleted with ageing. Proteases specific of or common to taproot and lateral roots were identified and the number of proteases corresponding to each category is indicated on top of each bar. The stars represent the senescence associated proteases (red stars) and the longevity associated proteases (blur stars). The number of cysteine, serine, aspartate metallo- and threonine proteases and of proteasome subunits associated to each categories is indicated in the table. Red colour intensify with increasing numbers.

A total of sixty-nine taproot specific proteases (41 over-accumulated and 28 depleted with senescence) were identified, while lateral root only contained sixteen (Figure 10). Amongst the forty-one senescence-related proteases of taproot, half are serine proteases, and we can note the presence of well-known senescence-related cysteine proteases as SAG12, RD21A and RD21B (Roberts et al. 2012). The taproot depleted proteases are mainly represented by proteasome subunits and serine proteases.

Five of the six cysteine protease inhibitors identified in the taproot and lateral root DAPs were more abundant in young root tissues (figure 11). By contrast, kunitz trypsin inhibitors that usually regulate serine protease activities, were more abundant in the old root tissues. The opposite senescence effect on the relative abundances of the different cysteine proteases and of their cognate inhibitors is in good accordance with a fine-tuned control of proteolytic activities along root development.

**Figure 11.**
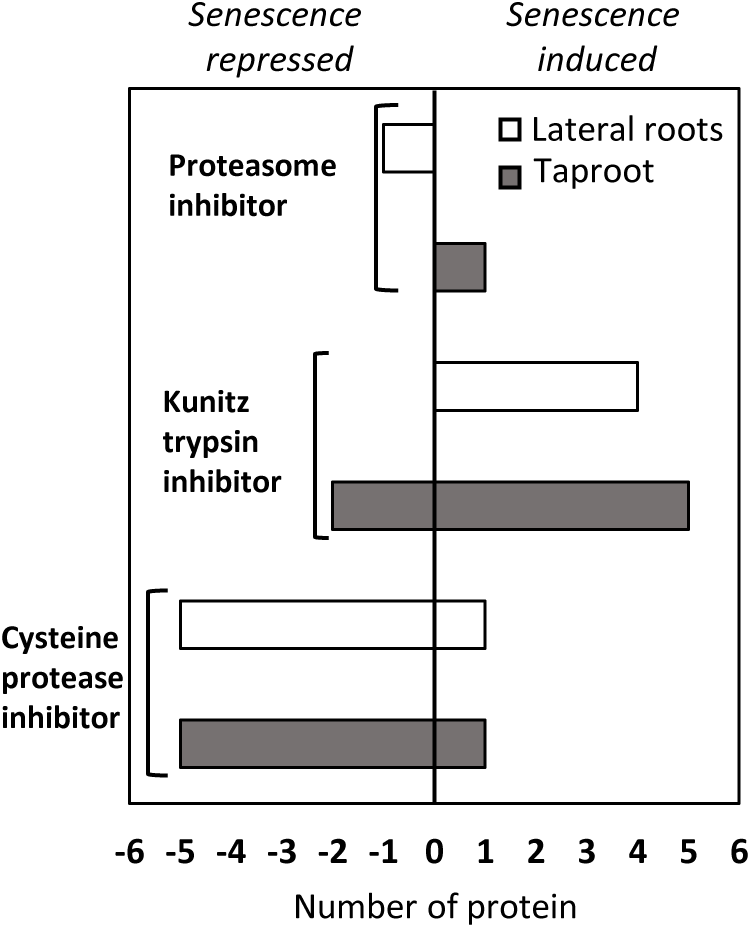
Evolution of protease inhibitors in taproot and lateral roots. Proteases inhibitors of taproot (grey bars) and lateral roots (white bars) were classified in senescence repressed or induced depending on if they were depleted or up-accumulated with ageing. The size of each bar in the histogram is associated with the number of protease inhibitors identified. Three main groups of proteases inhibitors were found, proteasome inhibitor, Kunitz trypsin inhibitor and cysteine protease inhibitor .

## DISCUSSION

This study focuses on the characterization of age-dependent changes occurring in the two distinct root compartments of *B. napus*, the taproot and the lateral roots. Plants were cultivated with sufficient nitrate (HN) and without nitrate (LN), which is a condition that has been previously reported as exacerbating the senescence and N remobilisation processes leading to nutrient transfer from the root to seeds for pod development in Arabidopsis (James et al., 2018).

### Anatomical evidences for root senescence

The three stages chosen for this study cover the entire development cycle of *B. napus*, and we can observe between T1 and T2 the classical drop in biomass for vegetative organs (Figure 2) as described previously (Malagoli et al., 2005; Girondé et al., 2015) that coincide with the formation of pods and of transfer of nutrients from remobilization of plant resources to the seed development. Browning of lateral roots and taproot is one of the phenotype of root ageing which appeared at T1 and was especially pronounced at T2 (Figure 1). As very few is known about root senescence, browning has been reported in several studies on grapes, peaches, populus or cotton as a trait characteristic of the shift from an active to a senescent root (Comas et al., 2000; Baldi et al., 2010; Wojciechowska et al., 2020; Zhu et al., 2021). Our light microscope observations confirmed the browning of root structure (Figure 3) is paralleled by cell deformation and by the loss of viability of the cortical cells in both the LN and HN plants. This loss of cell root integrity has been described as an anatomical trait named root cortical senescence (RCS) in several studies (Schneider et al., 2017; Schneider and Lynch, 2018; Liu et al., 2019; Lynch, 2019).

In contrast to lateral roots, no visual change in cell shape rwas observed time and whatever the N treatment suggesting different features for lateral root and taproot ageing (Figure 3-4). It can be observed that at the output of the vernalization period, taproot cells accumulated a large quantity of starch granules stored within numerous amyloplasts (Figure 4). Starch accumulation was independent of nitrate condition, in good agreement with Rossato et al., (2002a). Starch concentrations in the vernalized taproot represented 21.3 % and 22.3 % of the dry biomass in the HN and LN conditions, respectively. Starch concentration was 14 times higher than that measured in the lateral roots (Figure S3) confirming the importance of taproot as a carbon storage compartment. The dismantling of amyloplasts and the reduction of the size of starch granules observed in Figure 4 reflects the reallocation of carbon from the taproot to the above ground organs during the reproductive as sown by (Rossato et al., 2001; Rossato et al., 2002a). Ferritin deposits observed in the taproot amyloplasts, irrespective of nitrate conditions the N treatment increased at flowering (T1) and even more at pod development (T2) (Venzhik et al., 2019) were consistent with the accumulation of iron between the T1 and T2 (Figure 4 and S1). The proteomic data presented above confirm the higher abundance of ferritin proteins in taproot of LN plants (ferritin 1 (XP_013723422.1), 3 (XP_013648011.1, XP_022547543.1) and 4 (XP_013687876.1; Table S12). Overaccumulation of iron with taproot ageing could explain overproduction of reactive oxygen species resulting in taproot browning. Amyloplasts dismantling, which has already been reported in nongreening senescent cells (Inada et al., 2000) and the ferritin accumulation which is considered as a senescence marker in animals (Wiley and Campisi, 2021) and plant (Murgia et al., 2007) cells, show that taproot undergo senescence at T1 and T2 reproductive stages.

### Root proteins are N sources to sustain shoot growth and reproduction

Under HN, protein contents, amino acid contents and total N content evolve in parallel with aging in both taproot and lateral root (Figure 5). They increase from T0 to T1 reflecting nitrogen storage in the root compartments, and decrease from T1 to T2, reflecting the remobilization of N to the shoot. In contrast to HN, the increases in total N, proteins and amino acids at T1 was not observed under LN condition, but their decreases between T1 and T2 was significant in taproot and attested of N remobilization processes (Rossato et al., 2002b). In addition, the no accumulation of amino acids from proteolysis at T2 in either root organ amino acids supposed to be well exported, in line with rapeseed’s strong capacities to export amino acids (Tilsner et al., 2005). These results highlight the importance of the taproot and lateral roots, as N source organs especially during seed filling. Interestingly, soluble proteins are the main form of protein storage (figure 5C and 5D). Protein degradation in both root organ between T1 and T2 was consistent with the strong increase of protease activities at pH 5.5 and 7.5 at T2 in taproot under both LN and HN, and in lateral root when remobilization is enhanced by the shortage of nitrate under LN (Figure 6).

### Proteomic analysis of senescence markers in the lateral roots and taproot of LN plants

In order to identify changes in protein contents, to characterize senescence effects and to identify proteases involved in N remobilization in the taproot and in the lateral roots, a shotgun proteomic analysis was carried out on the roots of LN plants. Proteomes were largely modified by aging in both taproot and lateral roots, and gene ontology revealed that in both organs, processes related to catabolism and N metabolism were significantly modified.

The fine analysis of the classes of proteins modified in each organ exclusively, or common to taproot and lateral roots revealed interesting features (Figure 8). Specific taproot response to aging is reflected by the disappearance with ageing of protein categories related to nucleotide synthesis, translation, unfolded protein response (UPR) machinery that mitigates ER-stress, response to carbohydrates, and by the appearance with aging of protein categories related to amino-acid catabolism, protein quality control and cell-wall catabolism. Specific response of lateral roots is the disappearance with aging of anabolism categories related to fatty acids, amino-acids, organic acids syntheses and to sulfate assimilation, and the over-representation with aging of processes related to the disorganisation of protein complex, cytoskeletons, actin filaments, of alternative respiration, pentose phosphate related NADH production. The common proteins categories over-represented in the early taproot and lateral root development are linked to DNA replication, translation, redox control, translation, pyridine metabolism and risobome assembly, Ehrlish amino acid catabolism for the formation of aldehydes and alcohols and flavonoid synthesis. Proteins categories common to the senescent taproot and lateral roots include functions linked to the degradation of hydrogen peroxide, detoxification respiration and pentose metabolism. Thus globally, the SDP protein categories specific or common to taproot and lateral roots are mainly related to anabolic functions and synthesis of metabolites, and to cell growth and maintenance involving DNA replication, translation; the specific or common SAP protein categories are mainly related to catabolism of proteins, amino acids and cell wall, to detoxification and to alternative energy metabolisms. It can be noticed that our proteomic analysis provides a comprehensive picture of root senescence that globally recapitulates several senescence features described for leaves in many studies, with the exception that we do not find in root the aspects linked to chloroplast decay that are found in leaves (Breeze et al., 2011).

We then decided to try a comparison of the senescence-related changes in root protein contents in *B. napus* with ageing, with the changes recorded during leaf ageing in Arabidopsis for gene expressions (Breeze et al., 2011) (Figure 9). The exploration of *B. napus* proteins homologous to the senescence-repressed genes and senescence-associated genes reported by (Breeze et al., 2011) for Arabidopsis, led us to identify short lists of markers in common between leaf senescence and root senescence, in Arabidopsis and *B. napus* respectively. These markers include senescence-repressed and senescence-associated proteins that will facilitate further root senescence studies. In the senescence-repressed leaf and root markers, proteins related to C metabolism are over-represented in both taproot and lateral root, while in the senescence-associated leaf and root markers, proteins related to nucleobase catabolism and oxidoreductase are over-represented. The catabolism of nucleobases, and more particularly the catabolism of purine, could increase the pool of remobilisable nitrogen. In fact, nucleic acids and more specifically purine and its degradation products (glyoxylate and ammonia) could also be a significant source of nitrogen for the reproductive parts (Brychkova et al., 2008a; Brychkova et al., 2008b; Werner and Witte, 2011). In total, taproot and lateral roots share 75 senescence-repressed markers and 104 senescence associated markers that constitute a robust list of genes that characterize root and leaf senescence irrespective of root type.

### Protease changes with aging are different in taproot and lateral roots

Focusing more specifically on proteases which are a good marker of senescence, we found that a large set of proteases are modified in a similar way in taproot and lateral roots with ageing (Figure 10). These proteases include cysteine, serine and aspartate proteases that accumulate with ageing in both root types, while the young taproot and lateral root tissues are characterized by the fact they contain the same proteasome subunits and metallo-proteases (Figure 10).

Besides the proteases found in both taproot and lateral roots, a sticking feature is the fact that the taproot specific proteases were much more abundant than the lateral root specific proteases. While protease activities were globally lower in taproot than in lateral roots (Figure 6), it is surprising to see that the diversity of the proteases is higher in the taproot than in the lateral roots. As taproot is a storage organ, which is not the case of lateral root, we hypothesize that the taproot specific proteases are more specific of the degradation of the protein reserves and possibly of cell wall proteases as indicated by the gene ontology analysis. The senescence-associated proteases specifically found in taproot are mainly represented by serine carboxypeptidases and Clp protease subunits (Figure S4). Taproot accumulation of serine proteases such as SCPL7 (XP_013672786.1), SCPL28 (XP_013659509.1), SCPL29 (XP_013720411.1) and SCPL50 (XP_013674515.1), which are predicted as extracellular suggests their participation to cell wall modifications associated with senescence as reported in leaves by Borniego et al., (2020) and to programmed cell death as shown in the leaves of *Avena sativa* (Coffeen and Wolpert, 2004). The specific accumulation of ClpR1 (XP_013735849.1), ClpR2 (XP_013698293.1), ClpR4 (XP_013671352.1), Clp4 (XP_013670730.1) and Clp6 (XP_013657456.1), which are involved in chloroplast degradation during leaf senescence (Roberts et al., 2002; Kato and Sakamoto, 2010; Diaz-Mendoza et al., 2016), suggest their participation to the dismantling of amyloplasts observed taproot cells (Figure 4). It is interesting to note that the taproot specific aspartate proteases over-accumulated with senescence belong to the family A1 (Figure S4), like the CND41protease involved in chloroplast degradation during leaf senescence in *Nicotiana tabacum* and *Arabidopsis thaliana* (Kato et al., 2004; Diaz et al., 2008; James et al., 2018). All these aspartate proteases being predicted in the vacuolar and extracellular compartment, it cannot be excluded they could be involved also in cell wall modifications (Borniego et al., 2020).

The over-representation of cysteine proteases (CP) in senescing tissues is consistent with the picture of proteases changes in leaves with aging described in the literature (Buchanan-Wollaston and Ainsworth, 1996; Guo et al., 2004; Martínez et al., 2007; Roberts et al., 2012; Girondé et al., 2016; Poret et al., 2016) (Figure 10). These results are consistent with a repression of cysteine protease inhibitors in both root organs (Figure 11). CPs are known to be the class of proteases most overexpressed during natural or induced leaf senescence in numerous species and in particular rapeseed (Buchanan-Wollaston and Ainsworth, 1996; Guo et al., 2004; Martínez et al., 2007; Roberts et al., 2012; Girondé et al., 2016; Poret et al., 2016). Among CPs, Papain like cystein proteases (PLCPs) is the main subfamily associated with massive protein degradation during leaf senescence (Parrott et al., 2010) and some of them (SAG2, SAG12) are usually used as a standard marker for leaf senescence (Hensel et al., 1993; Noh and Amasino, 1999; Gombert et al., 2006). Several PLCPs which are found over-accumulated at the last stage especially in the taproot (RD21A, B, C, RD19A, C, Cathepsin B3, SAG12) are also found over-expressed in rapeseed or Arabidopsis leaves during natural or stress induced senescence (Noh and Amasino, 1999; Poret et al., 2016; Pružinská et al., 2017), making them excellent markers of root and leaf senescence (Figure S4).

## Conclusion

This study provides a comprehensive picture of the senescence related events occurring in taproot and lateral roots of *B. napus* with aging. This study identified senescence markers common to root and leaves and proteases potentially involved in taproot decay for N remobilization. The results presented paves the way for functional analyses of the senescence actors playing master roles in nutrient management and for the selection of new players facilitating the breeding of rapeseed varieties with improved taproot N storage and remobilization capacity at the root level.

## Acknowledgments

This work was supported by the French National Research Agency (ANR-19-CE14-0009-02 hAPPEN : Autophagy, Proteases and plant Performances). We are most grateful to PLATIN’ (Plateau d’Isotopie de Normandie) core facility for elemental analysis, Dr Didier Goux from CMABIO^3^ platform for the microscopy analyses. Dr Benoît Bernay from Proteogen platform for his help to proteomic analysis and Julie Frémont for her technical assistance. The authors declare that the research was conducted in the absence of any commercial or financial relationships that could be construed as a potential conflict of interest.

## Author Contributions

MJ, PE and JT conceived and designed the work. MJ conducted the experiments. MJ analyzed the data. TB performed the mass spectrometry analysis. MJ, PE, JT and CMD wrote the manuscript. AM and FC discussed and modified the work content. All authors carefully revised and approved the manuscript.

## Legends

**Figure S1. Iron concentration in taproot during the development of rapeseed cultivated under high (HN; A) or low nitrogen (LN; B) conditions.** The different harvest times T0, T1 and T2 correspond to vernalization output (B9), beginning of flowering (F1) and pod developments (G4) stages, respectively. Values are means ± SE (n=3). Significant differences between samples over time regardless the N condition are indicated by different letters (p-value ≤ 0.05).

**Figure S2. Clustering of the proteases modulated in the taproot and lateral root tissues during the development of rapeseed cultivated under low nitrogen (LN) conditions.** Proteases were selected using the MEROPS database and gene ontology. The different harvest times T0, T1 and T2 correspond to vernalization output (C1), beginning of flowering (F1) and pod developments (G4) stages, respectively. Heatmaps (A, B) represent the abundance of proteases (transformed in Z-score; red and blue for high and low accumulation, respectively) whose variability over time was confirmed by ANOVA in taproot (A) and in lateral roots (B). Each column corresponds to a biological replicate and the numbers at the left of the heatmap correspond to a cluster number.

**Figure S3. Starch content in lateral roots during the development of rapeseed cultivated under high (HN) or low nitrogen (LN) conditions.** The different harvest times T0, T1 and T2 correspond to vernalization output (B9), beginning of flowering (F1) and pod developments (G4) stages, respectively. Values are means ± SE (n=3). Significant differences between samples over time regardless the N condition are indicated by different letters (p-value ≤ 0.05).

**Figure S4. Proteases with significantly increased accumulation between beginning of flowering (T1) and pods development (T2) stages in taproot and lateral roots of rapeseed cultivated under low nitrogen (LN) condition.** Among proteases previously identified using the MEROPS database and gene ontology, only those significantly accumulated at least in one root organ between T1 and T2 are represented (FDR ≤ 0.05; n=3). The blue-red color gradient is proportional to the LogFC (from -2 to 2). The cluster numbers are those assigned in Figures 8A and 8C. Cysteine (yellow), aspartic (orange), serine (green), metallo-(grey), threonine protease (purple) and proteasome related proteins (blue) are abbreviated with Cys, Asp, Ser, Met, Thr and Prt, respectively. Predicted subcellular localizations of each protease, determined with the SUBA5 site, are cytoplasm (C), nucleus (N), endoplasmic reticulum (Er), extracellular (Ext), mitochondria (M), plastid (P), peroxisome (Px), and vacuole (V).

**Table S1. Identification and XIC-based quantification of proteins in the taproot under low nitrogen (LN) condition.** The Gene Onthology (GO) were provided by the panther database (pantherdb.org). Proteases identified by the MEROPS database and gene ontology are hylighted by a “Yes”. The different harvest times T0, T1 and T2 are associated to B9, F1 and G4 stages of rapeseed development, respectively.

**Table S2. Identification and spectral count (SC) quantification of proteins in the taproot under low nitrogen (LN) condition.** The Gene Onthology (GO) were provided by the panther database (pantherdb.org). Proteases identified by the MEROPS database and gene ontology are hylighted by a “Yes”. The different harvest times T0, T1 and T2 are associated to B9, F1 and G4 stages of rapeseed development, respectively.

**Table S3. Identification and XIC-based quantification of proteins in the lateral roots under low nitrogen (LN) condition.** The Gene Onthology (GO) were provided by the panther database (pantherdb.org). Proteases identified by the MEROPS database and gene ontology are hylighted by a “Yes”. The different harvest times T0, T1 and T2 are associated to B9, F1 and G4 stages of rapeseed development, respectively.

**Table S4. Identification and spectral count (SC) quantification of proteins in the lateral roots under low nitrogen (LN) condition.** The Gene Onthology (GO) were provided by the panther database (pantherdb.org). Proteases identified by the MEROPS database and gene ontology are hylighted by a “Yes”. The different harvest times T0, T1 and T2 are associated to B9, F1 and G4 stages of rapeseed development, respectively.

**Table S5. Identification of proteins in the taproot under low nitrogen (LN) condition in each cluster of the heatmap (**Figure 7A). Homologous genes of the B. napus senescence associated (SAP) and senescence depleted (SDP) proteins identified in taproot were identified amongst the senescence associated (SAG) and senescence repressed (SRG) genes from Breeze et al., (2011)

**Table S6. Identification of proteins in the lateral roots under low nitrogen (LN) condition in each cluster of the heatmap (**Figure 7B). Homologous genes of the B. napus senescence associated (SAP) and senescence depleted (SDP) proteins identified in lateral roots were identified amongst the senescence associated (SAG) and senescence repressed (SRG) genes from Breeze et al., (2011)

**Table S7. Biological processes enrichments in the taproot under low nitrogen (LN) conditions.** GO enrichment analyses were performed using Cytoscape (v3.9.1) plug-in ClueGO (v2.5.9) with the *Brassica napus* L. genome as a background. The biological process grouping is based on the cohen kappa coefficient use to define term-term interrelation and functional groups based on shared genes between the terms. The *Brassica napus* L. leaf senescent proteases identified by Poret et al,. (2019) are highlighted in the 29 most significant biological processes.

**Table S8. Biological processes enrichments in the lateral roots under low nitrogen (LN) conditions.** GO enrichment analyses were performed using Cytoscape (v3.9.1) plug-in ClueGO (v2.5.9) with the *Brassica napus* L. genome as a background. The biological process grouping is based on the cohen kappa coefficient use to define term-term interrelation and functional groups based on shared genes between the terms. The *Brassica napus* L. leaf senescent proteases identified by Poret et al,. (2019) are highlighted in the 29 most significant biological processes.

**Table S9. Biological processes enrichment of different clusters and their intersection under low nitrogen (LN) conditions.** Biological processes enrichment was performed with all proteins presented in each UpSet plot group using Panther (https://www.pantherdb.org/). Significant enrichment (Fischer test with FDR correction) are presented. According to (Figure 8A) and (Figure 8B) subsets used for Upset plots in (Figure 8C) are: T1, taproot cluster 1; T2, taproot cluster 2; T3, taproot cluster 3; T4, taproot cluster 4; L1, lateral roots cluster 1; L2, lateral roots cluster 2; L3, lateral roots cluster 3; L4, lateral roots cluster 4.

**Table S10. Identification of proteases in the taproot under low nitrogen (LN) condition in each cluster of the heatmap (**Figure 8C). Proteases identified by the MEROPS database and gene ontology.

**Table S11. Identification of proteases in the lateral roots under low nitrogen (LN) condition in each cluster of the heatmap (**Figure 8C). Proteases identified by the MEROPS database and gene ontology.

**Table S12. Abundance of proteins related to iron storage identified in the taproot under low nitrogen (LN) condition.** The different harvest times T0, T1 and T2 correspond to vernalization output (B9), beginning of flowering (F1) and pod developments (G4) stages, respectively. Proteins with significant variation of their abundance between two developmental stages have a LogFC value in bold (p-value ≤ 0.05; n = 3).

